# Resolving single-cell copy number profiling for large datasets

**DOI:** 10.1101/2022.02.09.479672

**Authors:** Ruohan Wang, Yuwei Zhang, Mengbo Wang, Xikang Feng, Jianping Wang, Shuai Cheng Li

## Abstract

The advances of single-cell DNA sequencing (scDNA-seq) enable us to characterize the genetic heterogeneity of cancer cells. However, the high noise and low coverage of scDNA-seq impede the estimation of copy number variations (CNVs). In addition, existing tools suffer from intensive execution time and often fail on large datasets. Here, we propose SeCNV, a novel method that leverages structural entropy, to profile the copy numbers. SeCNV adopts a *local* Gaussian kernel to construct a matrix, *depth congruent map*, capturing the similarities between any two bins along the genome. Then SeCNV partitions the genome into segments by minimizing the structural entropy from the depth congruent map. With the partition, SeCNV estimates the copy numbers within each segment for cells. We simulate nine datasets with various breakpoint distributions and amplitudes of noise to benchmark SeCNV. SeCNV achieves a robust performance, i.e., the F1-scores are higher than 0.95 for breakpoint detections, significantly outperforming state-of-the-art methods. SeCNV successfully processes large datasets (>50,000 cells) within four minutes while other tools failed to finish within the time limit, i.e., 120 hours. We apply SeCNV to single-nucleus sequencing (SNS) datasets from two breast cancer patients and acoustic cell tagmentation (ACT) sequencing datasets from eight breast cancer patients. SeCNV successfully reproduces the distinct subclones and infers tumor heterogeneity. SeCNV is available at https://github.com/deepomicslab/SeCNV.

## Introduction

Copy number variations (CNVs) are an important type of genomic variations^1^, which refer to the duplication or deletion of a kilobase-sized DNA fragment^2^. Somatic CNVs can amplify oncogenes or delete tumor suppressor genes^3^, which plays essential roles in the pathogenesis and metastases of cancers^4–6^. Accurate detection of CNVs helps prevent, prognosticate, and treat cancers^7^. The next-generation sequencing (NGS) technologies enable the comprehensive characterization of CNVs. However, bulk DNA sequencing gives an aggregation of copy-number profiles across cells, which immixed the heterogeneous cell populations in cancers^8^. Therefore, copy number profiling of single-cell DNA sequencing (scDNA-seq) provides the potential to identify subclones and resolve genetic heterogeneity in cancer evolution^9^.

Despite the significance of detecting CNVs from scDNA-seq data, the task is challenging because of the high noise of scDNA-seq^10^. The noises can come from multiple sources: (i) The read coverage of a single cell is low; that is, the noise ratio is much higher in scDNA-seq^11, 12^. (ii) The whole-genome amplification procedure propagates biases to read coverage^13^. (iii) Unlike bulk DNA sequencing, it is uneasy to identify negative samples and use them to correct bias^14^ in scDNA-seq. Therefore, the computational CNVs detection tools designed for bulk sequencing data are inapplicable for scDNA-seq samples^15^.

A method to profile single-cell copy numbers usually consists of four steps, alignment, normalization, segmentation, and absolute copy number calling^15, 16^. First, the reads are aligned or mapped to the reference genome. Potential biases affecting read coverage are corrected. Subsequently, segmentation, a significant step in copy number profiling^15^, is conducted. The genome is partitioned into contiguous segments in which copy numbers remain the same, and the segment boundaries are also referred to as breakpoints. Lastly, the shared copy number within each segment in each cell, the absolute copy number, is inferred.

Existing tools apply multiple approaches to normalize the read coverage and segment the genomes. During normalization, GC content and mappability are two well-investigated biases^16, 17^. For segmentation, most existing tools are based on statistical methods. HMMCopy^18^ applies hidden Markov models to model the CNVs along each chromosome. Ginkgo^10^ and SCNV^14^ use a circular binary segmentation (CBS) method^19^ with statistical testing to partition the genomes.

However, there remain several issues to be resolved. As aforementioned, most existing methods correct the biases caused by GC content and mappability in the normalization step. Some other types of biases are overlooked, like incorrect genome assembly in regions near centromeres and telomeres^20^, constraining accurate CNVs detection. In the segmentation step, many methods process each sample independently. Joint segmentation brings advantages to breakpoint detection, especially for scDNA-seq data. First, integrating the signals from shared breakpoints of different cells increases the sensitivity of breakpoint detection. Second, joint segmentation better utilizes the information across the samples from one batch, essential for single-cell cancer genomic analysis, like subclone identification. The methods based on objective function are more suitable for joint segmentation since the information from multiple samples can be collected in one function^15^. However, choosing suitable objective functions and efficient optimization algorithms are challenging. Copynumber^21^ applies the least-squares estimation for joint segmentation, but the performance is poor on simulated data^22^. Moreover, Copynumber excludes steps for normalization and absolute copy number calling, limiting its application. Recently, SCOPE^23^ is designed with a cross-sample segmentation approach, but the iterative segmentation procedure to optimize the Bayesian Information Criterion is time-consuming.

To address these issues, we propose an efficient tool, SeCNV, for single-cell copy number profiling. First, SeCNV identifies normal cells from the coefficient of variation (CV) density of cells. Using normal cells as negative controls, SeCNV infers the bin-specific bias, not limited to GC content and mappability, and normalizes the read count matrix. Next, SeCNV constructs the *depth congruent map* (DCM) for bins with a local Gaussian kernel function, making SeCNV focus on the most distinct read count differences for any two bins and skip the signals caused by noise. Then SeCNV optimizes the structural entropy^24^ of the DCM to group the bins into segments with the same copy number states. Finally, SeCNV calculates the absolute copy number in each segment and infers the copy number profiles for each cell. This work shows that SeCNV, which is based on structure entropy, is more efficient and robust to noise than the existing statistic-based methods.

## Results

### Overview of SeCNV

An overview of the SeCNV pipeline is shown in Figure 1. Taking scDNA-seq data as the input, the pipeline of SeCNV consists of four steps. First, the reads are aligned to the reference hg19 or hg38. After sorting and deduplication, consecutive bins are constructed along the genome. SeCNV generates a cell-bin read count matrix according to the number of reads mapped in each bin. Second, the normal cells are found with a density peak method and used as the negative control. Next, SeCNV calculates each bin’s bias, which may be caused by GC contents, mappability, and others. Then the normalized read count matrix is obtained. Third, SeCNV constructs the DCM for all the bins with a local Gaussian kernel function, then uses dynamic programming to minimize the structural entropy and finds the best segmentation. Forth, an integer approximation algorithm is applied to estimate each cell’s ploidy, and the integer copy number of each bin is determined. Finally, SeCNV generates the copy number profiles for all cells. The properties of SeCNV and other copy number profiling methods are compared in Table 1. SeCNV is available at https://github.com/deepomicslab/SeCNV. The output files of SeCNV can be uploaded to https://sc.deepomics.org/^25^ for visualization.

**Figure 1.**
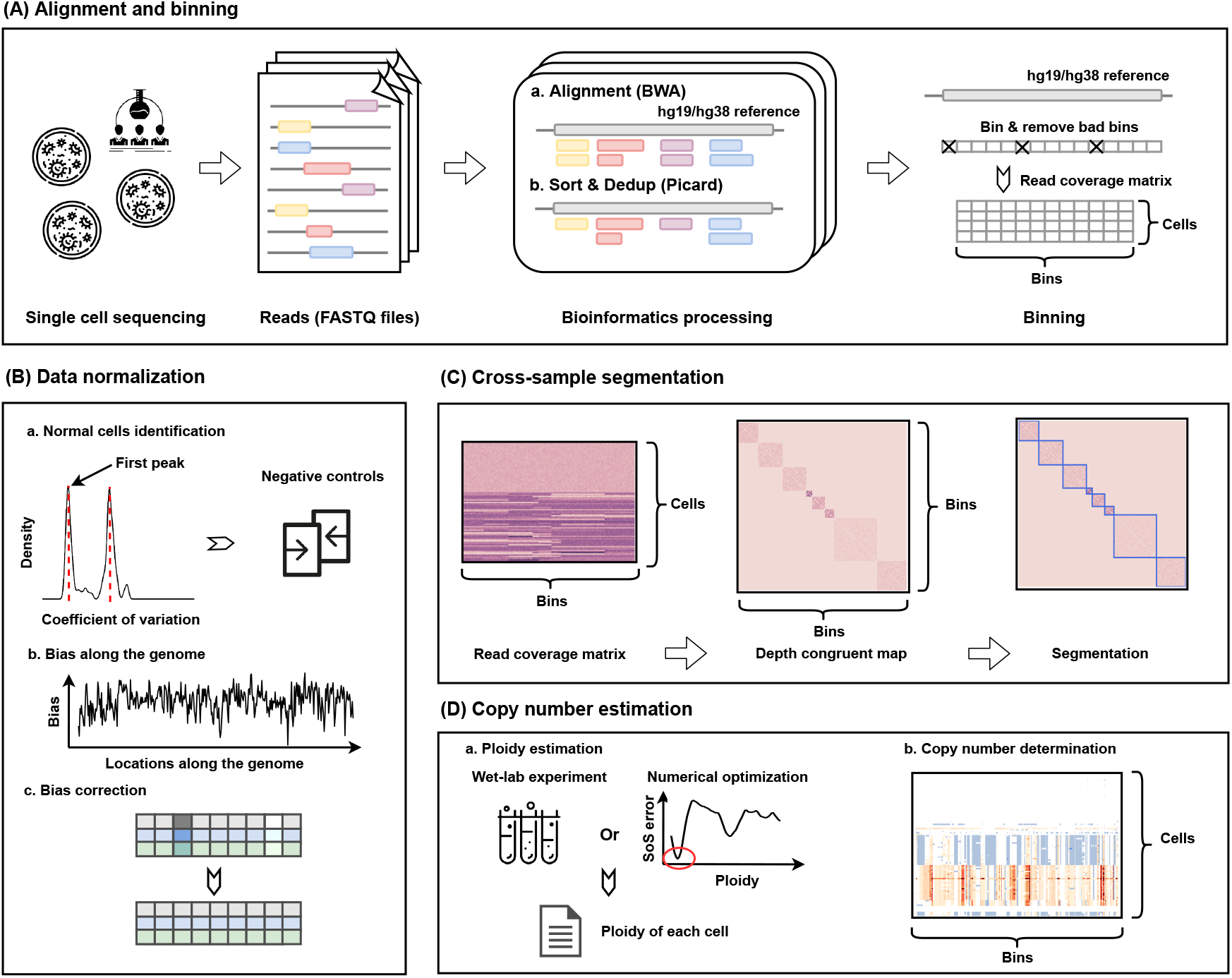
The overview of the SeCNV pipeline. The input of SeCNV is single-cell DNA sequencing data. The pipeline of SeCNV follows four steps to profile CNVs: A. Alignment and binning. B. Data normalization. C. Cross-sample segmentation. D. Copy number estimation. The output of SeCNV are the copy number profiles for all cells.

**Table 1.** Features of copy number profiling methods.

| Method | Alignment and binning | Normal cells identification | Bias correction | Segmentation | Copy number calling |
| --- | --- | --- | --- | --- | --- |
| **Copynumber**<sup>21</sup> | × | × | × | Cross-sample | × |
| **Ginkgo**<sup>10</sup> | Filtering outlier bins | × | GC correction | Single-sample | Integer approximation |
| **SCNV**<sup>14</sup> | Filtering outlier cells | With parameter | GC correction | Single-sample | Integer approximation |
| **SCICoNE**<sup>26</sup> | × | × | × | Cross-sample | Markov chain Monte Carlo |
| **SCOPE**<sup>23</sup> | Filtering outlier bins and cells | With parameter | GC and mappability correction | Cross-sample | Expectation-maximization |
| **SeCNV** | Filtering outlier bins and cells | Without parameter | Bin-specific correction | Cross-sample | Integer approximation |

### Evaluation of the segmentation procedure

To evaluate the segmentation performance of SeCNV, we compared SeCNV with the single-sample segmentation method CBS^19^ and the other three cross-sample segmentation methods, Copynumber^21^, SCICoNE^26^ and SCOPE^23^, on the nine simulated instances (see Methods). Since CBS is based on a single sample, we only count the breakpoints present in at least three samples. More parameter settings for the compared methods are shown in Table S1. We generated 100 instances for each case and calculated the average recall, precision, and F1-score values across the datasets for the five methods.

First, we evaluated the five methods on *Sim_norm_0.4, Sim_norm_0.6*, and *Sim_norm_0.8* (Figure 2 A-C, Table S2). In scDNA-seq, diploid cells are almost always collected together with aneuploid cells^20^. These normal cells are likely to weaken the breakpoint signals for cross-sample segmentation. When the percentage of normal cells increased from 40% to 80%, the average precision of CBS increased a lot since there were fewer false positives. The average recall of Copynumber decreased from 0.4060 to 0.0270. SCICoNE suffered from low recall on the three cases. SCOPE and SeCNV maintained a good performance on cases. The average recall and precision of SCOPE and SeCNV were higher than 0.9. Among the three methods, SeCNV obtained the highest average F1-scores, with recall and precision of almost 1, for all three cases.

**Figure 2.**
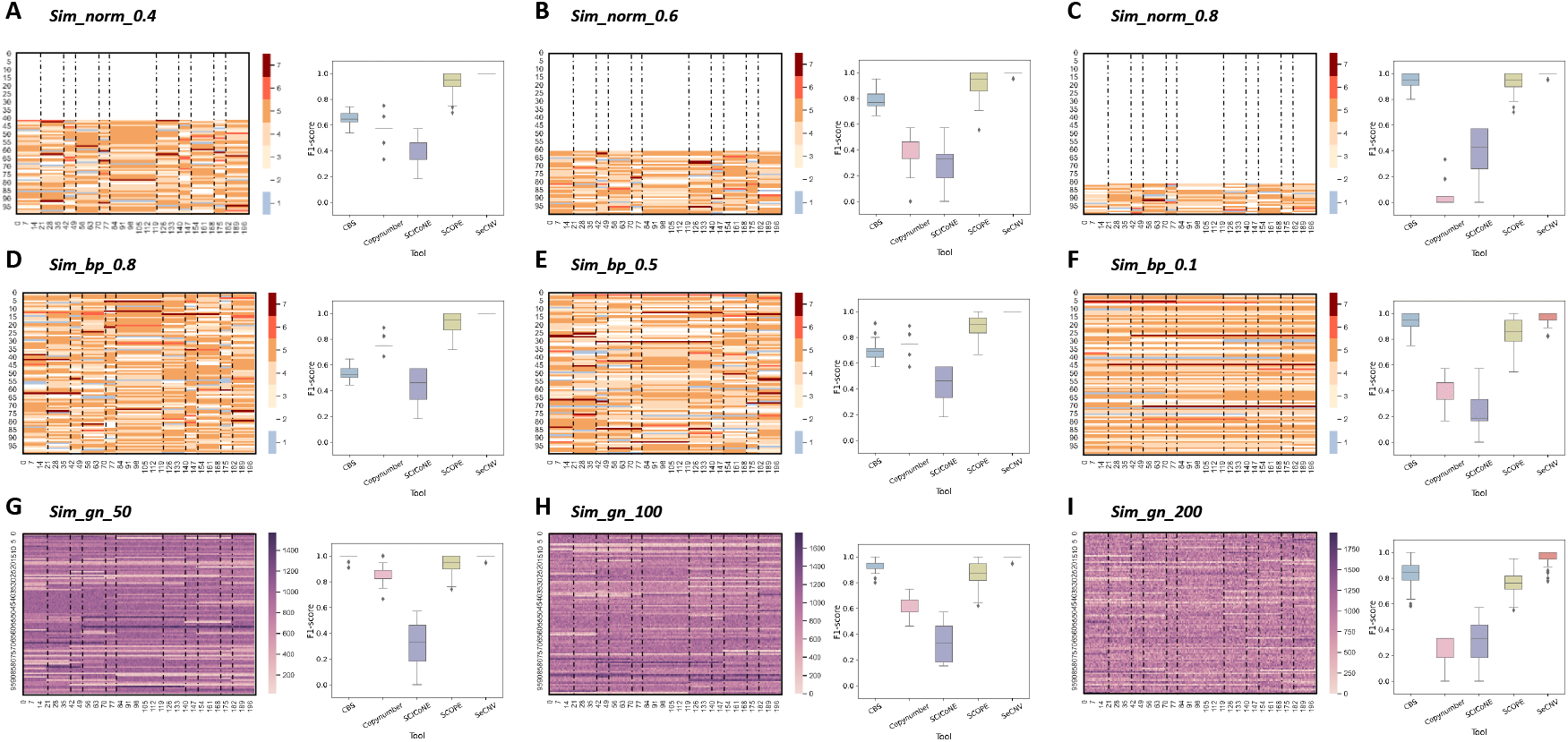
The comparison of segmentation performance on simulated datasets. We simulated three datasets: A. *Sim_norm_0.4* with 40% of the cells are normal cells. B. *Sim_norm_0.6* with 60% of the cells are normal cells. C. *Sim_norm_0.8* with 80% of the cells are normal cells. D. *Sim_bp_0.8* with breakpoint occurrence probability of 0.8. E. *Sim_bp_0.5* with breakpoint occurrence probability of 0.5. F. *Sim_bp_0.1* with breakpoint occurrence probability of 0.1. G. *Sim_gn_50* with Gaussian noise variance of 50. H. *Sim_gn_100* with Gaussian noise variance of 100. I. *Sim_gn_200* with Gaussian noise variance of 200. The left figures are the heatmaps of the copy numbers (A-F) or the read counts (G-I) for the 100 cells and 200 bins, and the right boxplots show the F1-scores of CBS, Copynumber, SCICoNE, SCOPE, and SeCNV on copy number segmentation.

Next, we applied the methods on *Sim_bp_0.8, Sim_bp_0.5*, and *Sim_bp_0.1* (Figure 2 D-F, Table S3) to check their performance on datasets with different breakpoint occurrence probabilities. Although the same genetic background cells usually share breakpoints, the breakpoints are unnecessary to appear in every cell from the same batch since different cells may come from various subclones. CBS obtained low precision when the breakpoint occurrence probability is high. SCICoNE obtained low recall across all the instances. The average recall of Copynumber decreased from 0.6180 to 0.2700, with the breakpoint occurrence probability decreased since the signal was diluted, while SCOPE and SeCNV kept a stable performance. SeCNV obtained the best average F1-scores, which are higher than 0.95, on all three cases.

Finally, we checked the influence of data noise on the segmentation performance of the five methods. In the simulated cases *Sim_gn_50, Sim_gn_100*, and *Sim_gn_200*, the variance of zero-mean Gaussian noises increased from 50 to 200 (Figure 2G-I, Table S4), which can be regarded as the noise amplitude. We noted that with the increase of Gaussian noise variance, the average recall of Copynumber and the average precision of CBS and SCOPE decreased by a wide margin while SeCNV kept a good performance on both metrics. For all three cases, SeCNV achieved the best F1-scores again.

To evaluate the efficiency of the segmentation procedure based on structural entropy, we compared the segmentation module of SeCNV with that of CBS, Copynumber, SCICoNE, and SCOPE on *in silico* datasets with different sizes. We simulated *Sim_norm_0.8, Sim_bp_0.1*, and *Sim_gn_200* datasets with 100 cells, 1,000 cells, 10,000 cells, 50,000 cells, and 100,000 cells. Then we compared the segmentation performance and CPU time for the five tools. As shown in Figure S1, SeCNV achieved higher F1 scores and consumed less CPU time on all cases. For large datasets (i.e., 50,000 cells or 100,000 cells), SeCNV successfully finished the segmentation process within four minutes while CBS, Copynumber, and SCICoNE suffered from inferior performance and SCOPE failed to yield results within the time limit (i.e., 120 hours).

### Analysis on single-nucleus sequencing data of breast cancer patients

We applied SeCNV to single-nucleus sequencing (SNS) data from two triple-negative breast cancer patients, genetically heterogeneous (polygenetic) T10 and genetically homogeneous (monogenetic) T16^27^ (Table S5). 100 single cells from T10 had been sequenced, and fluorescence-activated cell sorting (FACS) analysis suggested that there are four distributions of ploidy in T10: Diploids (D), Hypodiploids (H), Aneuploid A (AA), and Aneuploid B (AB). For T16, 52 cells from the primary tumor and 48 cells from its liver metastasis had been sequenced, and FACS showed two ploidy distributions: Diploids (D) and Aneuploids (A).

We set the bin length as 50 kb and used hg19 as the reference. SeCNV first identified the normal cells and used them as negative controls to normalize read counts for each cell. For illustration, we showed the normalization results for four cells from different subclones of the T10 dataset (Figure S2). The read counts were more stable along the genome after normalization for the diploid cells, as expected.

Figure 3 shows the estimated copy number results for T10 and T16, respectively. The heatmap of the estimated copy numbers for 99 cells from T10 (SRR054599 was removed because of the low coverage) shows obvious patterns for the four subclones. The heatmap of T16 (SRR089717 was removed because of the high CV value) shows similar copy number profiles between primary and metastatic tumor cells, indicating it is a monogenetic tumor type. The Pearson correlation coefficients between the ploidies from FACS analysis and the ploidies estimated by SeCNV were 0.9476 and 0.9563 for T10 and T16. Then we applied uniform manifold approximation and projection (UMAP)^28^ to copy numbers of all cells from T10 and T16 for dimensional reduction. We projected the copy number profiles to two dimensions for visualization. After dimensional reduction, four subclones in T10 were clearly separated, consistent with the FACS results. For T16, primary Aneuploids (PA) and metastatic Aneuploids (MA) separated despite the high similarity between the copy number profiles. All findings were highly concordant with previously reported results^27^. SeCNV also achieved good performance with different bin lengths and references (Section S1.1, Figure S3-S4).

**Figure 3:**
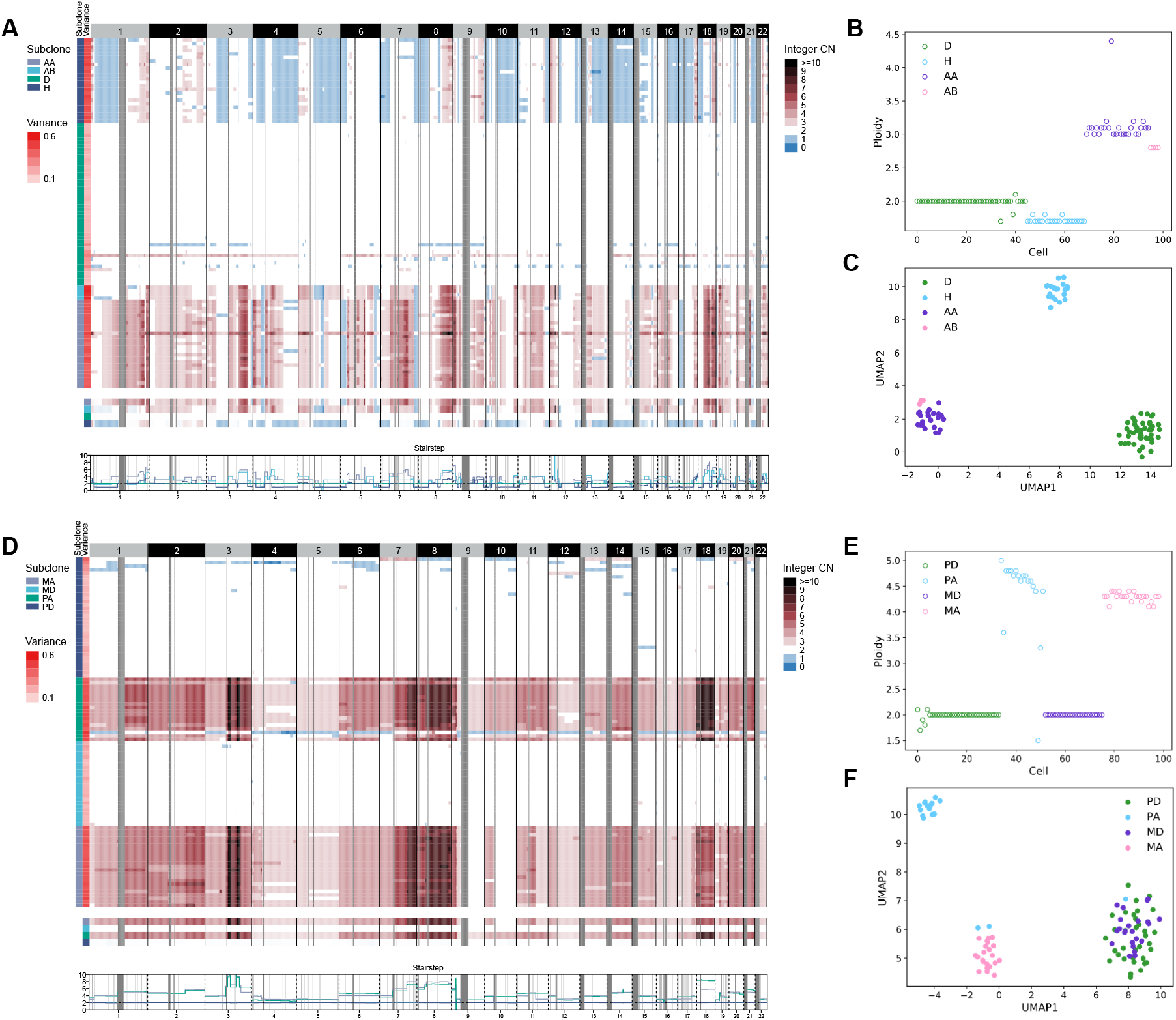
The copy number analysis of breast cancer patients T10 and T16 with SeCNV. A. The estimated copy number profiles of T10. The left columns show the subclones determined by FACS (Diploids (D), Hypodiploids (H), Aneuploid A (AA), and Aneuploid B (AB)) and CVs calculated by SeCNV. B. The ploidies of T10 cells generated by SeCNV. Cells from different subclones are shown in different colors. C. UMAP projections of the estimated copy number profiles of T10 cells. The cells from different subclones are apparently separated. D. The estimated copy number profiles of T16. The left columns show the subclones determined by FACS (primary Diploids (PD) and metastatic Diploids (MD), primary Aneuploids (PA), and metastatic Aneuploids (MA)) and CVs calculated by SeCNV. E. The ploidies of T16 cells generated by SeCNV. Cells from different subclones are shown in different colors. F. UMAP projections of the estimated copy number profiles of T16 cells. The cells from PD and MD fuse, and the cells from PA and MA are separated.

To further validate the CNVs called with SeCNV, we applied the copy number profiles from array comparative genomic hybridization (arrayCGH)^29^ of bulk samples as the ground truth. Since arrayCGH results for some subclones of T16 are unavailable, we only used the scDNA-seq data from T10 for evaluation. For the three non-diploid cell subclones from T10, we showed the copy numbers along the genome from arrayCGH experiments and SeCNV (Figure 4). Since CBS and Copynumber do not call absolute copy numbers, we excluded CBS and Copynumber in the experiment. Instead, we included Ginkgo^10^ in the comparison, which applied the CBS algorithm for segmentation. The lower panels in Figure 4 show the mean square error (MSE) between the estimated copy numbers and the ground truth for the three tools. SeCNV and SCOPE had a comparable performance on Hypodiploidy subclone and Hyperdiploidy AA subclone, whereas SeCNV achieved the best performance on Hyperdiploidy AB subclone. Ginkgo is a cloud-based analysis tool, so we compared the runtime of SCICoNE, SCOPE, and SeCNV on T10 and T16. The results are shown in Figure S5. SeCNV was five times faster than SCICoNE and seven times faster than SCOPE.

**Figure 4.**
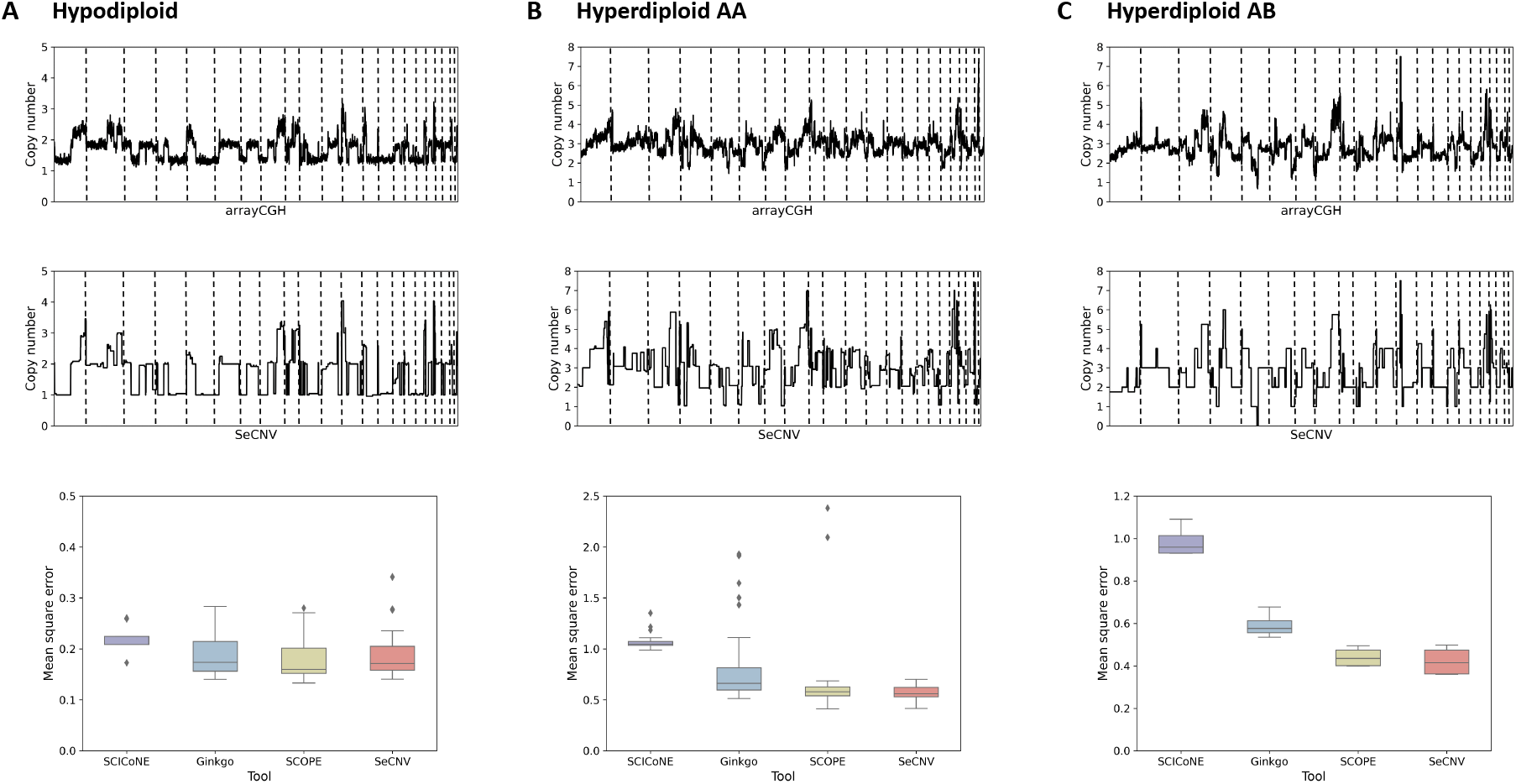
Comparison of the copy number profiling on T10 against CNV called from arrayCGH. For the three non-diploid cell subclones A. Hypodiploidy B. Hyperdiploidy AA C. Hyperdiploidy AB, the relative copy numbers from aGCH of bulk samples (upper) and SeCNV (middle) are compared. The dotted lines separate different chromosomes. The MSE between the copy number profiles obtained from aGCH and estimated from SCICoNE, Ginkgo, SCOPE, and SeCNV are compared in the boxplots (lower).

### Analysis of acoustic cell tagmentation sequencing data of breast cancer patients

We also utilized SeCNV to analyze recently published acoustic cell tagmentation (ACT) sequencing datasets from eight triple-negative breast cancer patients, each dataset containing ~1,000 cells (Table S5)^30^. First, we applied SeCNV to profile copy numbers for the eight datasets. We set the bin length as 50 kb and used hg19 as the reference. The ploidies estimated with SeCNV have similar distributions to the mean ploidies detected by FACS (Figure 5A). Additionally, SeCNV successfully reproduced the copy number profiles for the eight datasets (Figure 5B and Figure S6). The results show the cells from eight breast cancer patients are genetically homogeneous.

**Figure 5.**
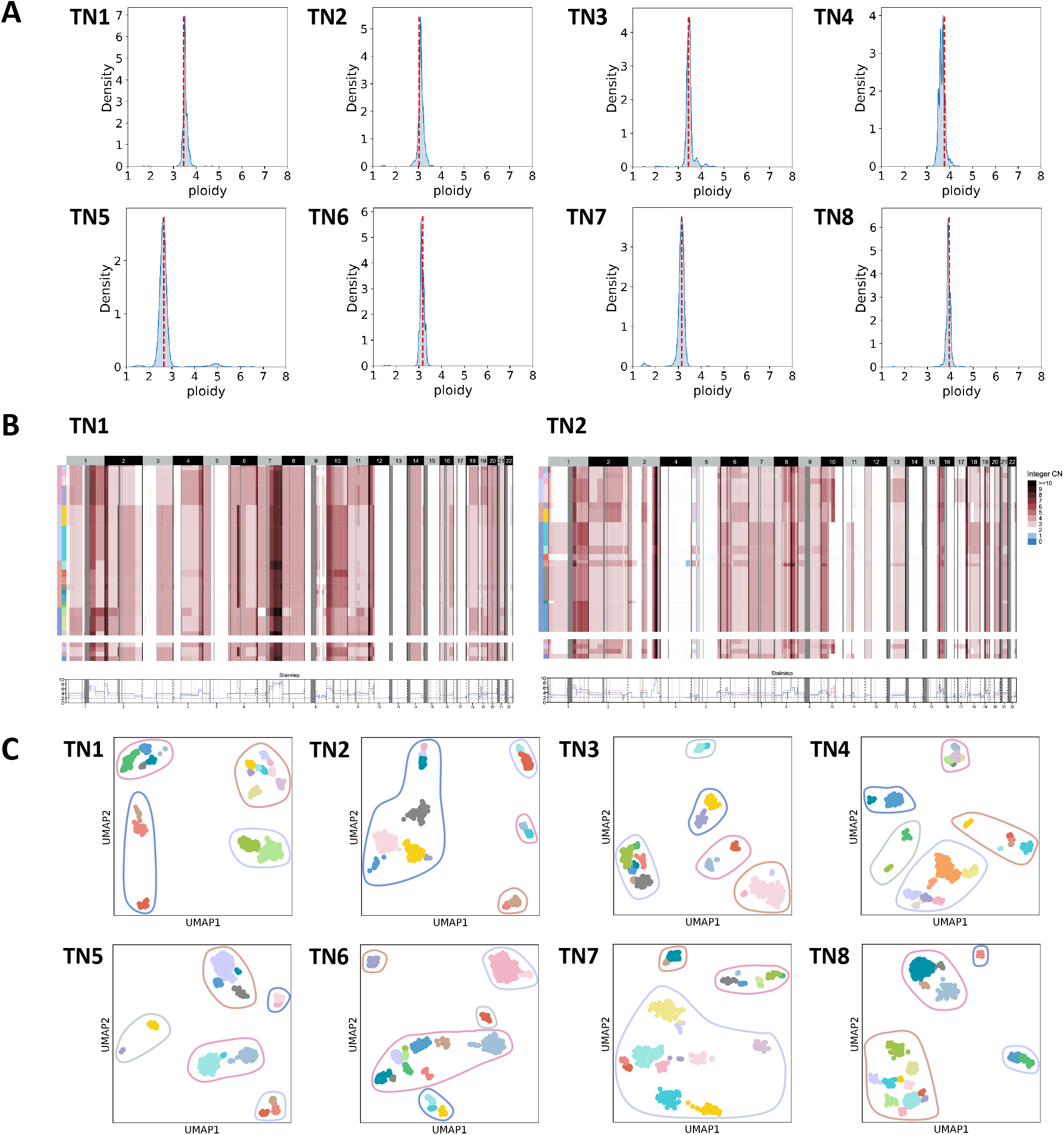
The ploidies, copy numbers, and clonal structures of the ACT sequencing data from breast cancer patients. A. The ploidy distributions of the eight patients estimated with SeCNV. The red lines show the mean ploidies detected by FACS. B. The copy number profiles of TN1 and TN2 estimated with SeCNV. The two left columns show the superclones and subclones determined with copy number clustering. C. UMAP projections of the estimated copy number profiles of the eight patients. Different colors show different subclones, and the circles separate different superclones.

Next, we identified superclones and subclones from the copy numbers called by SeCNV. The cells in the same clone should share similar copy number profiles. Following the previously reported steps^30^, we first constructed the shared nearest neighbor (SNN) graph for cells based on the called copy numbers. We identified the superclones as the connected subgraphs. For the cells in each superclones, we further classified them into subclones using DBSCAN^31^ on their UMAP embeddings. The parameter settings can be found in Table S6. Figure 5C shows the clonal structures for the eight datasets. Furthermore, we analyzed the breakpoint occurrence in different clones (Figure S7). We found that some breakpoints are shared across subclones, even superclones, while others only occur in unique subclones or superclones, which is concordant with previously analysis^30^. Cross-sample segmentation makes it easier to detect the shared breakpoints, thus holding great promise for identifying clonal structures.

## Discussion

SeCNV integrates binning, data normalization, cross-sample segmentation, and copy-number estimation to profile copy numbers for single cells. To overcome the low coverage and high noise of scDNA-seq, SeCNV applies a bin-specific normalization method instead of only considering the biases from GC content and mappability. Additionally, SeCNV applies a joint segmentation method to detect the shared breakpoints among the samples. Notably, In arrayCGH experiment, the mean of logarithmically transformed ratios is computed for color reversal experiments. The data are similar to the read count data from NGS experiments numerically^16^. Therefore, the segmentation algorithm can also be applied in arrayCGH data.

In the first step of the joint segmentation procedure, SeCNV constructs the DCM for bins along the whole genome with the normalized read counts. The values in the DCM present the similarities of copy number states between the bins. SeCNV uses a newly defined local Gaussian kernel function to capture the variation signals from high noise data. We have also tried other methods to construct the DCM, and it turns out the local Gaussian kernel function achieves the best performance (Section S1.2, Table S7-S11). After obtaining the DCM, SeCNV applies a dynamic programming algorithm to optimize structural entropy. The segmentation with the lowest structural entropy is returned as the optimal solution. Structural entropy measures the dynamic complexity of graphs in structural information theory, and minimizing structural entropy is equivalent to deciphering graph structure. Therefore, structural entropy can be used as the objective function for DCM segmentation. Recently, some researchers applied structural entropy to detect topologically associated domains in Hi-C data and achieved good performance^32–34^. In this work, we utilized structural entropy on copy number segmentation. We believe structural entropy can be applied to various segmentation and clustering problems in biology.

Based on the properties of scDNA-seq, we simulate nine datasets to benchmark SeCNV. In the simulated cases *Sim_norm_0.4, Sim_norm_0.6, Sim_norm_0.8, Sim_bp_0.8, Sim_bp_0.5*, and *Sim_bp_0.1*, the presence of normal cells and small subclones weaken the breakpoint signals. In comparison, SeCNV is more sensitive to these cases. Copynumber fails to capture the signals when the breakpoint occurrence probability is small, which is concordant with previously reported results^15, 22^. The cases *Sim_gn_50, Sim_gn_100*, and *Sim_gn_200* simulate the high noises of scDNA-seq, SeCNV achieves the highest F1-score on the three cases, indicating that our structural-entropy-based method exhibits an extraordinary ability to tolerant noise compared with statistic-based methods.

We apply SeCNV on scDNA-seq datasets from breast cancer patients with different sequencing protocols. With FACS and arrayCGH data as the ground truth, SeCNV successfully normalizes data, infers the ploidies, and estimates the copy numbers along the genome. Then we reproduce the distinct subclones with the inferred copy number profiles and decipher the tumor heterogeneity. The findings are highly consistent with FACS and arrayCGH results. Models of tumor progression at the cellular level have been studied for a long time. With the improvement of scDNA-seq technologies and copy number profiling methods, it is likely to infer the tumor evolutionary history from the knowledge of genetic heterogeneity.

## Materials and methods

### Alignment and binning

Taking single-cell DNA sequencing data as input, SeCNV aligns reads to the reference genome with BWA^35^ firstly. Then, SeCNV applies the bioinformatic preprocessing pipeline of SCOPE^23^. In detail, Picard^36^ is applied to sort bam files, add read groups, and remove duplicates. Next, SeCNV constructs genome-wide consecutive bins with a user-defined bin size. We download the 100-mers mappability track for hg19 reference from the ENCODE Project (http://hgdownload.cse.ucsc.edu) and convert it to hg38 coordinates with UCSC liftOver tool. Following the CNV calling convention, we partition the genome into bins of a fixed number of nucleic acids. SeCNV computes each bin’s average GC content and mappability and masks the bins with extreme GC content or mappability. After eliminating the reads mapped to multiple genomic locations, SeCNV adopts Bedtools to place the reads to bins and generates the cell-bin read count matrix. Cells with mapped reads proportion (the ratio of the number of reads mapped to reference to the number of all sequencing reads) less than 0.2 are removed for quality control^23^. We denote the raw read count matrix as 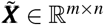, where *m* is the number of cells, and *n* is the number of bins. For notation simplicity, we use the index *c* to represent cells and the index *b* to represent bins, where 1 ≤ *c* ≤ *m* and 1 ≤ *b* ≤ *n*.

### Data normalization

After obtaining the read count matrix, SeCNV corrects for the bias of each bin and generates the normalzied read count matrix 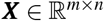 from 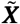. The normalization procedure is shown in Figure 6. First, SeCNV calculates the CV of read count in all bins for each cell. The CV value^37^ for a cell *c* calculated as

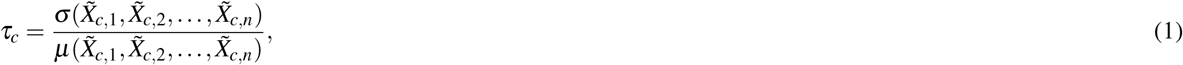

where *σ* is the standard deviation and *μ* is the mean of data.

After obtaining the set of CV values {*τ*_1_,…, *τ_m_*}, the Gaussian kernel density is calculated as

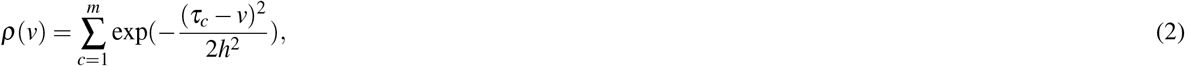

where *ρ* (*v*) denotes the density estimated at CV value *v* and *h* denotes the bandwidth parameter.

Since the cells are heterogeneous, the CV density curve is expected to have multiple peaks. Among all cells, the normal cells should have the lowest CVs, leading to the first peak. SeCNV detects the first peak by finding the lowest *v*\* satisfies that *ρ*(*v*\*) ≥ *ρ*(*v*) for each *v* in [*v*\* – *ε*, *v*\* + *ε*], for some *ε* > 0. SeCNV assigns the cells with CV values in the first density peak as normal cells.

**Figure 6.**
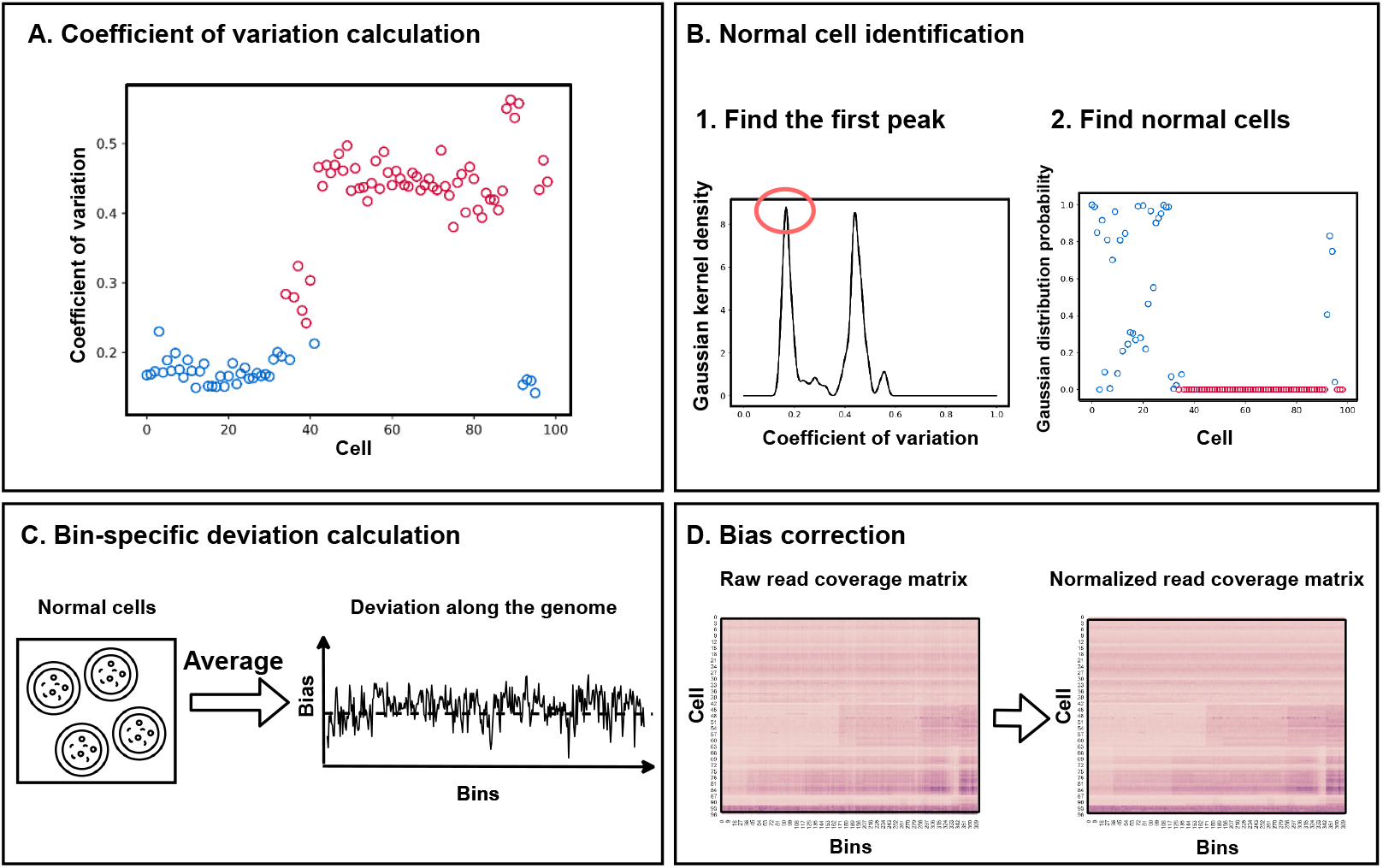
The data normalization procedure. A. First, SeCNV calculates the CV values for all the cells. B. Then SeCNV estimates the Gaussian kernel density for CV values and uses the first Gaussian kernel function to find the normal cells. C.Using the identified normal cells as the negative control, SeCNV calculates the bin-specific deviation along the genome. D. Finally, the read counts for all the cells are corrected.

Denote the identified normal cells set as ***v*** ⊂ {1,…,*m*}, where *m* is the number of cells. SeCNV regards these cells as negative control samples, which should have uniform count along the genome. Then we can determine the deviation of count from the genome median. For any normal cell *c* ∈ ***v***, SeCNV calculates the deviation for each bin *b* as 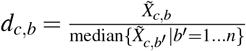, where *n* is the number of bins. Then the averaged deviations among all the normal cells are computed as 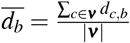. Finally, for each cell *j*, the biases are corrected with 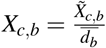.

### Cross-sample segmentation

The cross-sample segmentation step partitions the bins into non-overlapping segments of shared copy numbers. SeCNV applies a cross-sample segmentation procedure to identify breakpoints that are shared across cells. Figure 7 illustrates our proposed cross-sample segmentation algorithm.

**Figure 7.**
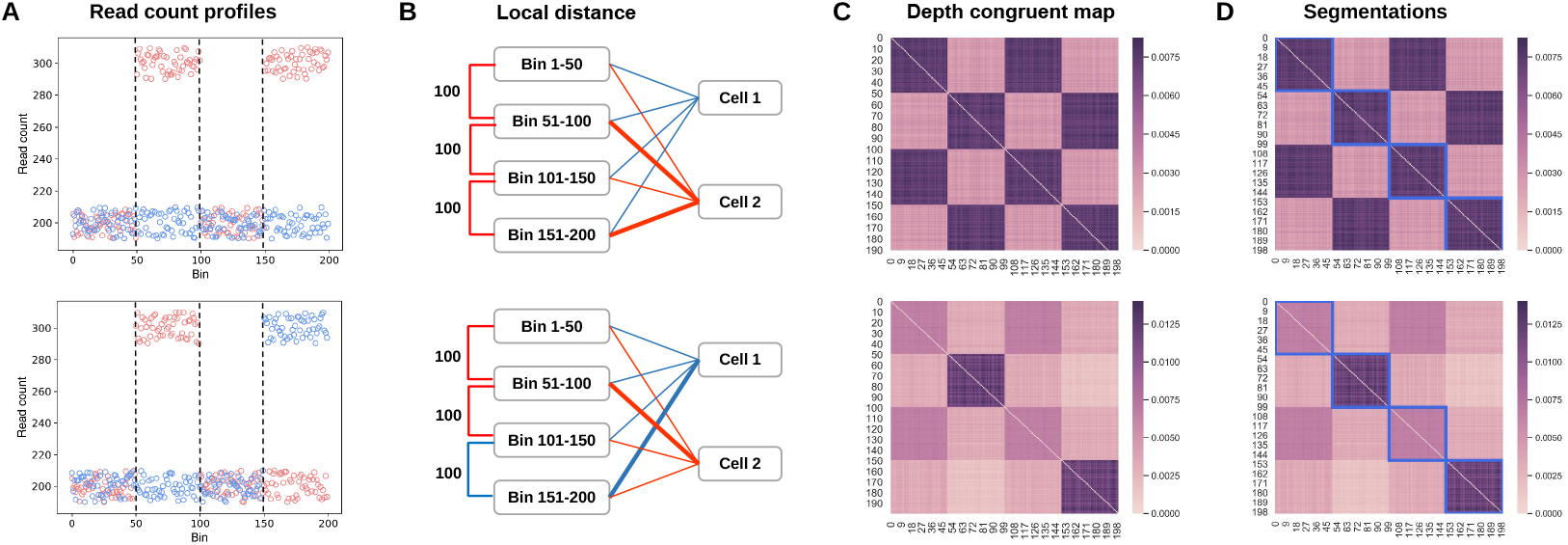
The cross-sample segmentation procedure. A. We generate two synthetic cases, each containing two cells (shown in red and blue respectively) and 200 bins with breakpoints located at 50, 100, and 150. The copy numbers alternate between 2 and 3, making the read counts alternate between 200 and 300. B. The local *L_2_* distances between neighboring bins (K=1). The width of lines between bins and cells indicates the read count values of the bin in the cell. C. The heatmap of the depth congruent map for bins. The intensity of the color can represent the weight of the edge between two bins. D. The segmentation results produced by SeCNV. SeCNV correctly finds the breakpoints at 50, 100, and 150.

First, we construct the DCM 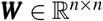 for all bins from the read count matrix 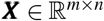, where *m* is the number of cells and n is the number of bins. ***W*** is a symmetric matrix, where the larger value of entry ***W***_*b,b*′_ indicating it is more likely that some cells have the same depth (or copy number) in bins *b* and *b*′. To preserve the copy number variations over the noise, we introduce a local *L*_2_ Gaussian kernel function which selects *K* cells with the most distinct differences for each pair of bins. Suppose *b* and *b*′ are two bins, then for *b* and *b*′, the read counts for all cells are {*X*_1,b_,*X*_2,b_,…,*X*_*m*,*b*_} and {*X*_1,b′_,*X*_2,*b*_′,…,*X*_*m,b*′_}, respectively. We denote ***z*** = {*z*_1_,*z*_2_,…,*z_m_*}, where *z_c_* = (*X*_*c,b*_–*X*_*c,b*′_)^2^. Then we choose a permutation *γ*: *V* → *V* that sorts ***z***, so that *Z*_*γ*(1)_ ≥ *Z*_*γ*(2)_ ≥… ≥ *Z*_*γ*(m)_. The local *L*_2_ distance between *b* and *b*′ can be computed as 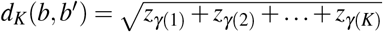. Finally, we use the local Gaussian kernel to compute the similarities between any two bins along the genome, leading to the DCM ***W*** with 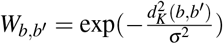. We set *K* = 5 and 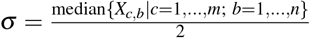 as defaults. All experiments in this paper were conducted with the defaults.

Next, SeCNV adopts structural entropy as the objective function to separate each genome sequence into segments homogeneous in copy number. A DCM is equivalent to an undirected graph, where each vertex represents a bin. Then segmenting the bin set into segments is equivalent to partitioning the vertex set into non-overlapping modules. Suppose that *P* = {*M*_1_,…, *M_j_*,…, *M_k_*}, 1 <*j* < *k*, is a partition of bin set along the genome, where *M*_1_ = {*b*_1_*l*__,…, *b*_1_*r*__}, *M_j_* = {*b_j_l__*,…, *b_j_r__*}, *M_k_* = {*b_k_l__*,…, *b_k_r__*}, *b_j_l__*, ≤ *b_j_r__*. Clearly, *b*_1_*l*__ = 1, *b_k_r__* = *n*, and *b_j_r__* + 1 = *b*_*j*_+1_*l*___. Then structural entropy on the partition *P* for the DCM ***W***is then defined as follows

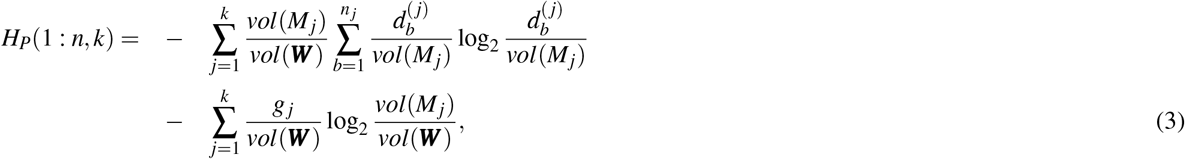

where 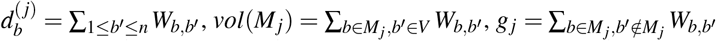, and *vol*(***W***) = ∑_1≤*b,b*′≤n_*W*_*b,b*′_.

We want to find a partition with minimal structural entropy for the DCM. Denote the optimal structural entropy of a DCM with *n* bins and *k* modules as

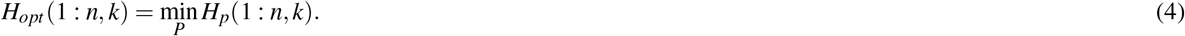

The optimal structural entropy of the DCM ***W*** is then defined as

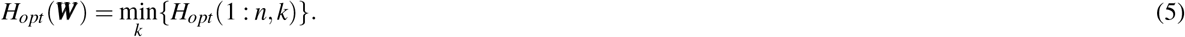

We define the structural entropy of the module containing the bins from *b* to *b*′ as *S*(*b*: *b*′), *b* ≤ *b*′. Then we can have the recurrent equation

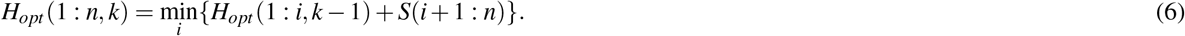

The equation enables us to find the optimal segmentation with dynamic programming for *k* partition. The algorithm is shown in Algorithm 1. The input is the DCM. First, SeCNV computes the dynamic programming table Γ and index table Ψ. Γ(*b, k*) records the optimal structure entropy for partition the first *b* vertices into *k* modules. Ψ(*b, k*) records the location of *k*-th breakpoint for the optimal segmentation. Then SeCNV chooses the *k* value with the optimal structure entropy as the final number of breakpoints and tracks each breakpoint’s locations with two tables. Finally, SeCNV reports the optimal segmentation.

#### Algorithm 1 Optimize structural entropy

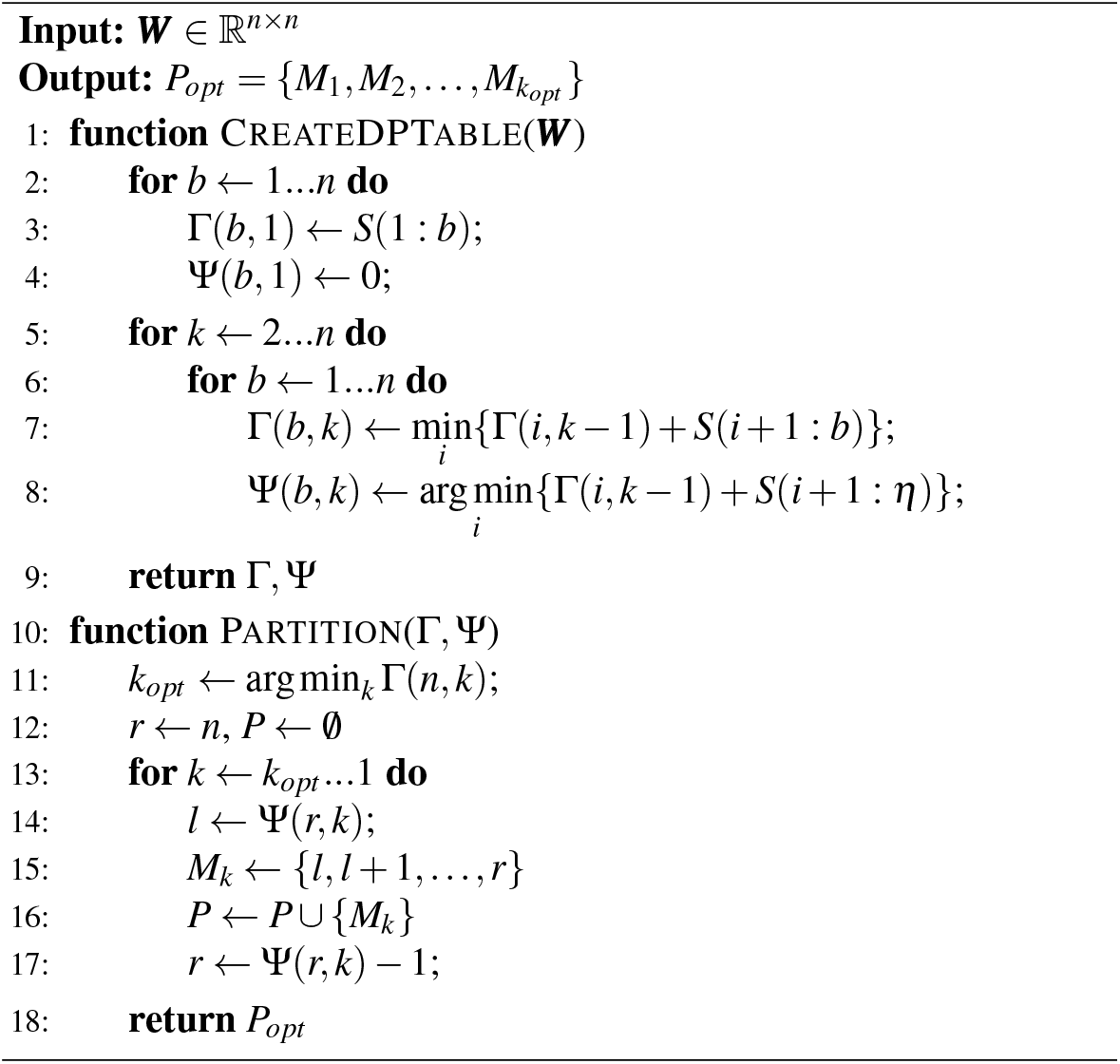

After segmentation, SeCNV replaces the read count in all bins of each segment with the median read count in the segment.

### Copy-number estimation

After segmentation, SeCNV can determine the copy-number state of each sample with the information of ploidy, which is the average of the estimated copy number across the genome. In some wet lab experiments, each cell’s ploidy can be determined directly by fluorescence with flow cytometry. When the ploidy information is unavailable from the scDNA-seq experiment, SeCNV applies an integer approximation algorithm^10, 14^. Since the copy numbers should be integers, this algorithm enumerates all the possible ploidy, for example, [1.50, 1.55, 1.60,…, 4.95,5.00]. Then it chooses the ploidy that minimizes the sum-of-squares error between the scaled copy numbers and their rounded integer copy numbers. Finally, SeCNV reports the rounded integer copy numbers as the final copy numbers.

### Simulated datasets

We simulated datasets with CNVs to evaluate the segmentation performance of SeCNV. All of the simulated datasets included 100 cells, and the genome of each cell was divided into 200 bins. Then we created ten breakpoints as the ground truth and simulated some CNV events on some randomly chosen cells while others were set to be normal cells. The absolute copy number within each segment varied from one to seven. We designed the Poisson distribution Pois(200c) to model the read counts for each bin, where *c* is the absolute copy number. To simulate the high noises of scDNA-seq, we added zero-mean Gaussian noise 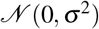 with different variances σ (i.e. standard deviation) to read counts. We designed nine cases to test the performance of SeCNV under different situations (Table S12). The normal cell percentage denotes the ratio of the number of cells without CNVs to the total number of cells; the breakpoint occurrence probability denotes the probability of breakpoints appearing in a cell; the Gaussian noise variance denotes the variance of zero-mean Gaussian noise added to the read counts.

## Acknowledgements

This work was supported by Strategic Interdisciplinary Research Grant (7005215). We would like to thank Dr. CHEN Lingxi, from the Department of Computer Science, City University of Hong Kong, for helping us do the visualization.

## Author contributions

RW designed the methods, performed most experiments, and drafted the manuscript. YZ and MW helped performing the experiments. XF collected the data. JW supervised this project and revised the manuscript. SCL proposed the topic, supervised this project, and revised the manuscript. All authors have read and approved of the final manuscript.

