## Supplemental notes, tables, and figures for "Resolving single-cell copy number profiling for large datasets"

### Supplementary Materials

#### S1 Supplementary Notes

##### S1.1 Hyper-parameter evaluation

First, we show the influence of the hyper-parameter $K$ for the Gaussian kernel function. The choice of $K$ affects how many cells are considered to construct the DCM for bins. In the manuscript, we recommend setting $K$ as 5 since the setting has the most robust performance. Here, we show the performance of SeCNV when $K$ is set to be 50 (Table S6) and 100 (Table S7) on the simulated datasets. We found SeCNV with large $K$ keeps good performance on most cases except for the dataset with high noise (*Sim_gn_200*). Smaller $K$ forces the model to focus on the most distinct variations instead of all the variations, including the noise.

Next, we evaluate the performance of SeCNV with different bin lengths and references. We set bin lengths to be 10 kb, 20 kb, 40 kb, 50 kb, 60 kb, and 80 kb, then check the performance of SeCNV on T10 and T16 datasets. For evaluation, we calculated the Pearson correlation coefficients between the ploidies from FACS analysis and the ploidies estimated by SeCNV. As shown in Figure S2, SeCNV achieves a robust performance with different bin lengths. Additionally, we use hg38 as the reference for SeCNV. SeCNV successfully reproduces the copy number profiles for T10 and T16 (Figure S3). The Pearson correlation coefficients between the ploidies from FACS analysis and the ploidies estimated by SeCNV are 0.9476 and 0.9236 for T10 and T16, respectively.

##### S1.2 Depth congruent map construction method comparison

Besides the local Gaussian kernel, we also tried three others methods to construct the depth congruent map (DCM) for bins. They are the Pearson-coefficient-based method, random-walk-based method, and Riemannian-manifold-based method. Our purpose is to construct the DCM $\boldsymbol{W}$ of shape $n * n$ for all the bins from the $m * n$ read depth matrix, where $m$ is the number of cells, and $n$ is the number of bins.

###### S1.2.1 Pearson-coefficient-based method

Pearson correlation coefficient is a widely-used measurement of the correlation between data sets. The Pearson correlation coefficient can be calculated as:

$$COV(b, b') = \frac{1}{m-1} \sum_{i=1}^{m} (b_i - \bar{b})(b'_i - \bar{b'}) \tag{S1}$$

In our problem, we use the value of the Pearson correlation coefficient to measure the similarity between bins. Large values indicate higher similarity. Negative values are set to be zero. Therefore, the DCM can be computed as $W_{b,b'} = max(0, COV(b, b'))$.

The performance of the Pearson-coefficient-based method on simulated datasets is shown in Table S8. Despite the high precision, the method obtains a low recall, especially when the subclone is small and the noise is high.

##### S1.2.2 Random-walk-based method

Firstly, we regard the read depth matrix as a $n*m$ bipartite graph, where n is the number of cells and m is the number of bins. The vertices can be divided into two disjoint sets *primary nodes* (bins) and *feature nodes* (cells). A edge connect a vertex in *primary nodes* with a vertex in *feature nodes* denotes the read count for the bin of the cell. Then our goal is to project the bipartite graph into a unipartite graph which represents the similarity between each bin. Consider a discrete time Markov process on the bipartite graph with $n$ *primary nodes* and $m$ *feature nodes*:

$$\begin{aligned} \boldsymbol{p}_p &= \boldsymbol{X} D_f^{-1} \boldsymbol{p}_f \\ \boldsymbol{p}_f &= \boldsymbol{X}^T D_p^{-1} \boldsymbol{p}_p \end{aligned} \tag{S2}$$

where $\boldsymbol{p}_p$ is the $n*1$ *primary nodes* probability vector, $\boldsymbol{p}_f$ is the $m*1$ *feature nodes* probability vector, $D_p$ is the diagonal matrix containing the degree of each *primary nodes*, $D_f$ is the diagonal matrix containing the degree of each *feature nodes*, and $X$ is the $n*m$ read depth matrix.

If a random walker start at a *primary nodes* and take two steps on the graph, then the walker will also stop at a *primary nodes*. We have the $n*1$ *primary nodes* probability vector after two steps:

$$\boldsymbol{p}_p = \boldsymbol{X} D_f^{-1} \boldsymbol{X}^T D_p^{-1} \boldsymbol{p}_p \tag{S3}$$

Clearly $AD_f^{-1}A^T D_p^{-1}$ is the transfer probability after two steps between *primary nodes*. Next we can built a symmetric matrix, which indicates the virtual edges between *primary nodes*, with the following equation:

$$\boldsymbol{W'} = D_p(AD_f^{-1}A^T D_p^{-1})^T \tag{S4}$$

If *primary nodes* $i$ is similar with *primary nodes* $j$, the three virtual edges $W'_{i,i}$, $W'_{i,j}$, and $W'_{j,j}$ should be close to each other. Based on this observation, we can construct the DCM for all the *primary nodes*:

$$W_{b,b'} = \frac{K(W'_{b,b'}, W'_{b,b}) + K(W'_{b,b'}, W'_{b',b'})}{2} \tag{S5}$$

where $K(W'_{b,b'}, W'_{b,b}) = e^{-\frac{d^2(w'_{b,b'}, w'_{b,b})}{\sigma^2_{b'}}}$ and $K(W'_{b,b'}, W'_{b',b'}) = e^{-\frac{d^2(w'_{b,b'}, w'_{b',b'})}{\sigma^2_{b}}}$ are the Gaussian kernel functions with local scaling.

The performance of the random-walk-based method on the simulated datasets is shown in Table S9. When the subclone is small and noise is high, the recall and precision of the random-walk-based method decrease by a wide margin since the breakpoint signals are diluted.

##### S1.2.3 Riemannian-manifold-based method

The advanced dimension redcution algorithm Uniform Manifold Approximation and Projection (UMAP) uses a weighted k-neighbour graph to represent a manifold with the following equation:

$$W'_{b,b'} = \exp(-\frac{-max(0, d(b,b') - \rho_b)}{\sigma_b}) \tag{S6}$$

Based on the local-connectivity constraint, $\rho_b$ and $\sigma_b$ can be defined as:

$$\begin{aligned} &\rho_b = \min\{d(b,b') | 1 \leq b' \leq k\} \\ &\sum_{b'=1}^{k} \exp(\frac{-\max(0, d(b,b') - \rho_b)}{\sigma_b}) = \log_2(k) \end{aligned} \tag{S7}$$

We use Euclidean distance in equation S7. Then the symmetric DCM can be computed as:

$$W_{b,b'} = W'_{b,b'} + {W'_{b,b'}}^T - W'_{b,b'} \cdot {W'_{b,b'}}^T \tag{S8}$$

The performance of the Riemannian-manifold-based method on the simulated datasets is shown in Table S10. The Riemannian-manifold-based method achieves good recall in most cases, demonstrating its ability to preserve the global topological structure of data. However, the precision is low, especially when noise is high.

### S2 Supplementary Tables and Figures

Table S1: **The parameters of the methods used for comparisons.**

| Method | Implementation | Function and parameters |
|---|---|---|
| **CBS** | R package | segment(smoothed.CNA.object, verbose=1) |
| **Copynumber** | R package | multipcf(data=data.wins,verbose=FALSE) |
| **SCICoNE** | Python package | sci.detect_breakpoints(chr_data, threshold=2.0);<br>sci.learn_tree(chr_data, bps['segmented_region_sizes'], full=False, n_reps=4, max_tries=4) |
| **SCOPE** | R package | normalize_scope_foreach(Y_qc=Y, gc_qc=ref$gc,K=1, ploidyInt=ploidy.sim,<br>norm_index=which(Gini¡=0.12), T=1:7,beta0=beta.hat.noK.sim, nCores=2);<br>segment_CBScs(Y=Y, Yhat=Yhat, sampname=colnames(Y), ref=ref,<br>chr=chri, mod =”integer”, max.ns=1) |
| **Ginkgo** | Website | Genome:Human (hg 19) |

Table S2: **The average recall, precision, and F1-score of CBS, Copynumber, SCICoNE, SCOPE, and SeCNV on copy number segmentation of simulated datasets with different normal cell percentages, the best F1-scores are shown in bold.**

| | **CBS** | | | **Copynumber** | | | **SCICoNE** | | |
|---|---|---|---|---|---|---|---|---|---|
| | Recall | Precision | F1-score | Recall | Precision | F1-score | Recall | Precision | F1-score |
| *Sim_norm_0.4* | 1 | 0.4827 | 0.6493 | 0.4060 | 1 | 0.5745 | 0.2280 | 1 | 0.3634 |
| *Sim_norm_0.6* | 1 | 0.6566 | 0.7896 | 0.2160 | 0.9600 | 0.3461 | 0.1970 | 0.9832 | 0.3188 |
| *Sim_norm_0.8* | 1 | 0.9180 | 0.9552 | 0.0270 | 0.2500 | 0.0485 | 0.2480 | 0.8792 | 0.3743 |

| | **SCOPE** | | | **SeCNV** | | |
|---|---|---|---|---|---|---|
| | Recall | Precision | F1-score | Recall | Precision | F1-score |
| *Sim_norm_0.4* | 0.9580 | 0.9249 | 0.9395 | 1 | 1 | **1** |
| *Sim_norm_0.6* | 0.9430 | 0.9000 | 0.9186 | 1 | 0.9982 | **0.9990** |
| *Sim_norm_0.8* | 0.9500 | 0.9145 | 0.9297 | 1 | 0.9900 | **0.9948** |

Table S3: **The average recall, precision, and F1-score of CBS, Copynumber, SCICoNE, SCOPE, and SeCNV on copy number segmentation of simulated datasets with different breakpoint occurrence probabilities, the best F1-scores are shown in bold.**

| | **CBS** | | | **Copynumber** | | | **SCICoNE** | | |
|---|---|---|---|---|---|---|---|---|---|
| | Recall | Precision | F1-score | Recall | Precision | F1-score | Recall | Precision | F1-score |
| *Sim_bp_0.8* | 1 | 0.3679 | 0.5365 | 0.6180 | 1 | 0.7631 | 0.2740 | 1 | 0.4209 |
| *Sim_bp_0.5* | 1 | 0.5251 | 0.6856 | 0.6030 | 1 | 0.7510 | 0.2960 | 1 | 0.4465 |
| *Sim_bp_0.1* | 0.9120 | 0.9685 | 0.9368 | 0.2700 | 0.9725 | 0.4184 | 0.1490 | 0.7212 | 0.2379 |

| | **SCOPE** | | | **SeCNV** | | |
|---|---|---|---|---|---|---|
| | Recall | Precision | F1-score | Recall | Precision | F1-score |
| *Sim_bp_0.8* | 0.9630 | 0.8937 | 0.9245 | 1 | 1 | **1** |
| *Sim_bp_0.5* | 0.9700 | 0.8308 | 0.8897 | 1 | 1 | **1** |
| *Sim_bp_0.1* | 0.9190 | 0.7901 | 0.8431 | 0.9220 | 1 | **0.9577** |

Table S4: **The average recall, precision, and F1-score of CBS, Copynumber, SCICoNE, SCOPE, and SeCNV on copy number segmentation of simulated datasets with different Gaussian noises, the best F1-scores are shown in bold.**

| | **CBS** | | | **Copynumber** | | | **SCICoNE** | | |
|---|---|---|---|---|---|---|---|---|---|
| | Recall | Precision | F1-score | Recall | Precision | F1-score | Recall | Precision | F1-score |
| *Sim_gn_50* | 1 | 0.9867 | 0.9929 | 0.7380 | 1 | 0.8454 | 0.2330 | 0.9312 | 0.3604 |
| *Sim_gn_100* | 1 | 0.8867 | 0.9379 | 0.4290 | 1 | 0.5975 | 0.2260 | 0.9137 | 0.3491 |
| *Sim_gn_200* | 0.8970 | 0.7782 | 0.8289 | 0.1650 | 0.9200 | 0.2779 | 0.2060 | 0.8068 | 0.3161 |

| | **SCOPE** | | | **SeCNV** | | |
|---|---|---|---|---|---|---|
| | Recall | Precision | F1-score | Recall | Precision | F1-score |
| *Sim_gn_50* | 0.9540 | 0.9288 | 0.9390 | 0.9980 | 1 | **0.9989** |
| *Sim_gn_100* | 0.9440 | 0.8130 | 0.8671 | 0.9960 | 1 | **0.9979** |
| *Sim_gn_200* | 0.9080 | 0.6526 | 0.7545 | 0.9610 | 0.9702 | **0.9648** |

Table S5: **Single-cell DNA sequencing datasets adopted in the study.**

| ScDNA-seq Protocol | Patient | Tumour type | Number of cells | Ploidy |
|---|---|---|---|---|
| DOP-PCR | *T10* | TNBC | 100 | 1.7/3/3.3 |
| | *T16* | TNBC | 100 | 4.1 |
| Acoustic cell tagmentation | *TN1* | TNBC | 900 | 3.45 |
| | *TN2* | TNBC | 824 | 3.03 |
| | *TN3* | TNBC | 1101 | 3.44 |
| | *TN4* | TNBC | 1307 | 3.76 |
| | *TN5* | TNBC | 1238 | 2.65 |
| | *TN6* | TNBC | 1205 | 3.17 |
| | *TN7* | TNBC | 907 | 3.15 |
| | *TN8* | TNBC | 1224 | 3.95 |

(TNBC:Triple-negative breast cancer)

Table S6: **Superclone and subclone identification parameters (k: the number of nearest-neighbor to construct the SNN graph; min_samples: the number of samples in a neighborhood for a point to be considered as a core point).**

| Patients | SNN graph | DBSCAN |
|---|---|---|
| *TN1* | k=100 | min_samples=12 |
| *TN2* | k=100 | min_samples=8 |
| *TN3* | k=100 | min_samples=6 |
| *TN4* | k=150 | min_samples=10 |
| *TN5* | k=100 | min_samples=8 |
| *TN6* | k=66 | min_samples=8 |
| *TN7* | k=100 | min_samples=8 |
| *TN8* | k=100 | min_samples=8 |

Table S7: **The average recall, precision, F1-score, and runtime of Gaussian kernel function with local distance (K=50) on copy number segmentation of simulated datasets.**

| | Recall | Precision | F1-score | Runtime (seconds) |
|---|---|---|---|---|
| *Sim_norm_0.4* | 1 | 0.9742 | 0.9864 | 754 |
| *Sim_norm_0.6* | 0.9840 | 0.7509 | 0.8497 | 747 |
| *Sim_norm_0.8* | 0.9600 | 0.7186 | 0.8203 | 733 |

| | Recall | Precision | F1-score | Runtime (seconds) |
|---|---|---|---|---|
| *Sim_bp_0.8* | 1 | 1 | 1 | 727 |
| *Sim_bp_0.5* | 1 | 1 | 1 | 740 |
| *Sim_bp_0.1* | 0.9590 | 0.9970 | 0.9768 | 729 |

| | Recall | Precision | F1-score | Runtime (seconds) |
|---|---|---|---|---|
| *Sim_gn_50* | 1 | 1 | 1 | 732 |
| *Sim_gn_100* | 1 | 0.9982 | 0.9990 | 760 |
| *Sim_gn_200* | 0.8880 | 0.6679 | 0.7607 | 737 |

Table S8: **The average recall, precision, F1-score, and runtime of Gaussian kernel function with local distance (K=100) on copy number segmentation of simulated datasets.**

| | Recall | Precision | F1-score | Runtime (seconds) |
|---|---|---|---|---|
| *Sim_norm_0.4* | 1 | 0.9689 | 0.9835 | 892 |
| *Sim_norm_0.6* | 0.9810 | 0.7436 | 0.8444 | 759 |
| *Sim_norm_0.8* | 0.9520 | 0.7073 | 0.8102 | 822 |

| | Recall | Precision | F1-score | Runtime (seconds) |
|---|---|---|---|---|
| *Sim_bp_0.8* | 1 | 0.9991 | 0.9995 | 873 |
| *Sim_bp_0.5* | 1 | 1 | 1 | 867 |
| *Sim_bp_0.1* | 0.9600 | 0.9943 | 0.9759 | 859 |

| | Recall | Precision | F1-score | Runtime (seconds) |
|---|---|---|---|---|
| *Sim_gn_50* | 1 | 1 | 1 | 863 |
| *Sim_gn_100* | 1 | 0.9982 | 0.9990 | 850 |
| *Sim_gn_200* | 0.8770 | 0.6583 | 0.7506 | 802 |

Table S9: **The average recall, precision, F1-score, and runtime of Pearson-coefficient-based method on copy number segmentation of simulated datasets.**

| | Recall | Precision | F1-score | Runtime (seconds) |
|---|---|---|---|---|
| *Sim_norm_0.4* | 0.3200 | 0.9967 | 0.4803 | 1091 |
| *Sim_norm_0.6* | 0.2200 | 0.9600 | 0.3557 | 1074 |
| *Sim_norm_0.8* | 0.1910 | 0.8850 | 0.3129 | 1076 |

| | Recall | Precision | F1-score | Runtime (seconds) |
|---|---|---|---|---|
| *Sim_bp_0.8* | 0.7582 | 0.8242 | 0.7887 | 1125 |
| *Sim_bp_0.5* | 0.4720 | 0.9700 | 0.6327 | 1100 |
| *Sim_bp_0.1* | 0.1600 | 0.7850 | 0.2655 | 1041 |

| | Recall | Precision | F1-score | Runtime (seconds) |
|---|---|---|---|---|
| *Sim_gn_50* | 0.2670 | 0.9767 | 0.4169 | 1075 |
| *Sim_gn_100* | 0.2680 | 0.9850 | 0.4190 | 1077 |
| *Sim_gn_200* | 0.2510 | 0.9100 | 0.3911 | 1071 |

Table S10: **The average recall, precision, F1-score, and runtime of random-walk-based method on copy number segmentation of simulated datasets.**

| | Recall | Precision | F1-score | Runtime (seconds) |
|---|---|---|---|---|
| *Sim_norm_0.4* | 0.9580 | 0.9506 | 0.9528 | 898 |
| *Sim_norm_0.6* | 0.9100 | 0.8626 | 0.8823 | 818 |
| *Sim_norm_0.8* | 0.7390 | 0.7426 | 0.7354 | 766 |

| | Recall | Precision | F1-score | Runtime (seconds) |
|---|---|---|---|---|
| *Sim_bp_0.8* | 0.9850 | 0.9846 | 0.9845 | 877 |
| *Sim_bp_0.5* | 0.9110 | 0.9555 | 0.9310 | 874 |
| *Sim_bp_0.1* | 0.4040 | 0.6391 | 0.4899 | 853 |

| | Recall | Precision | F1-score | Runtime (seconds) |
|---|---|---|---|---|
| *Sim_gn_50* | 0.7420 | 0.8605 | 0.7931 | 875 |
| *Sim_gn_100* | 0.6650 | 0.8289 | 0.7340 | 798 |
| *Sim_gn_200* | 0.5520 | 0.8440 | 0.6635 | 765 |

Table S11: **The average recall, precision, F1-score, and runtime of Riemannian-manifold-based method on copy number segmentation of simulated datasets.**

| | Recall | Precision | F1-score | Runtime (seconds) |
|---|---|---|---|---|
| *Sim_norm_0.4* | 1 | 0.7981 | 0.8848 | 754 |
| *Sim_norm_0.6* | 0.998 | 0.7932 | 0.8817 | 747 |
| *Sim_norm_0.8* | 0.9990 | 0.7945 | 0.8825 | 733 |

| | Recall | Precision | F1-score | Runtime (seconds) |
|---|---|---|---|---|
| *Sim_bp_0.8* | 0.9980 | 0.7890 | 0.8784 | 727 |
| *Sim_bp_0.5* | 0.9960 | 0.7889 | 0.8774 | 740 |
| *Sim_bp_0.1* | 0.9450 | 0.7360 | 0.8249 | 729 |

| | Recall | Precision | F1-score | Runtime (seconds) |
|---|---|---|---|---|
| *Sim_gn_50* | 0.9960 | 0.7811 | 0.8730 | 732 |
| *Sim_gn_100* | 0.9950 | 0.7810 | 0.8727 | 760 |
| *Sim_gn_200* | 0.9190 | 0.7133 | 0.8013 | 737 |

Table S12: **The parameters of the nine simulated cases.**

| Case | Normal cell percentage | Breakpoint occurrence probability | Gaussian noise variance |
|---|---|---|---|
| *Sim_norm_0.4* | 40% | 100% | 100 |
| *Sim_norm_0.6* | 60% | 100% | 100 |
| *Sim_norm_0.8* | 80% | 100% | 100 |
| *Sim_bp_0.8* | 0% | 80% | 100 |
| *Sim_bp_0.5* | 0% | 50% | 100 |
| *Sim_bp_0.1* | 0% | 10% | 100 |
| *Sim_gn_50* | 0% | 20% | 50 |
| *Sim_gn_100* | 0% | 20% | 100 |
| *Sim_gn_200* | 0% | 20% | 200 |

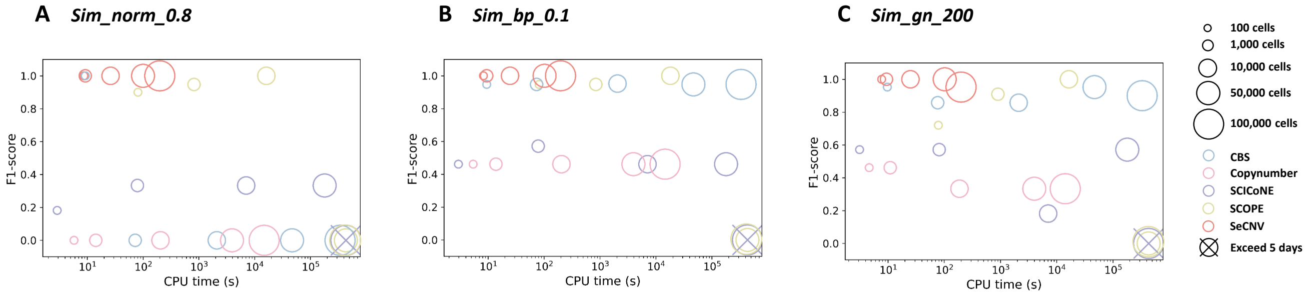


Figure S1: **CPU time and F1-score comparison on datasets with different size.** For Copynumber, SCOPE, and SeCNV, we compare their CPU time and F1-score on datasets with 100 cells, 1,000 cells, 10,000 cells, 50,000 cells, and 100,000 cells. Methods that exceed 5 CPU days limit are killed.

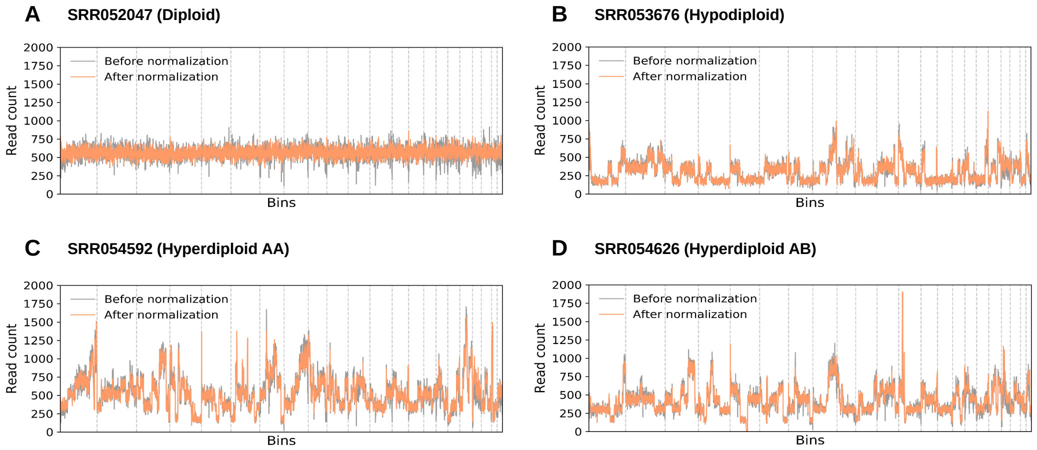


Figure S2: **SeCNV normalizes the read counts for scDNA-seq data from breast cancer patients T10.** For four cells from different subclones of T10, (A) a diploid cell, (B) a Hypodiploid cell, (C) a Hyperdiploid AA cell, and (D) a Hyperdiploid AB cell, the read counts before normalization and after normalization are shown. The dotted lines separate different chromosomes.

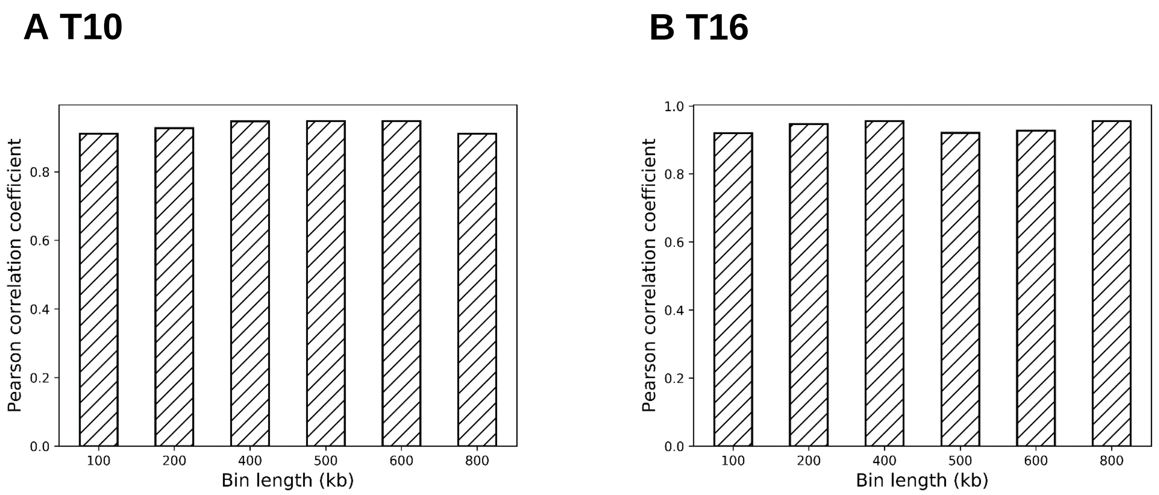


Figure S3: **The Pearson correlation coefficients between the ploidies from ground truth and the ploidies estimated with SeCNV of different size of bin.** For cells from T10 (A) and cells from T16 (B), the performance of SeCNV is robust with different length of bin from 100 kb to 800 kb.

**A T10**

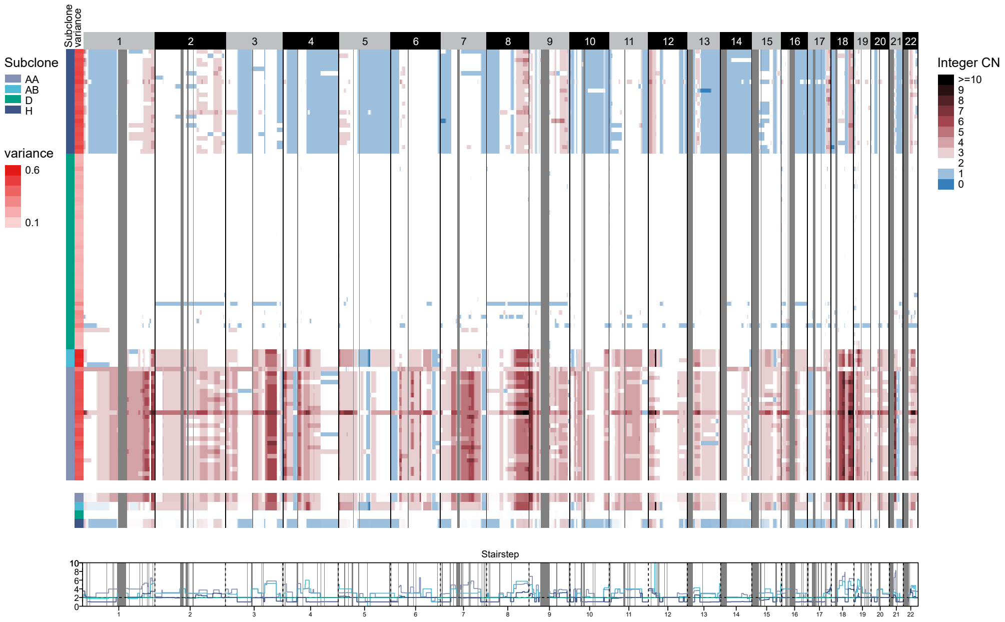


**B T16**

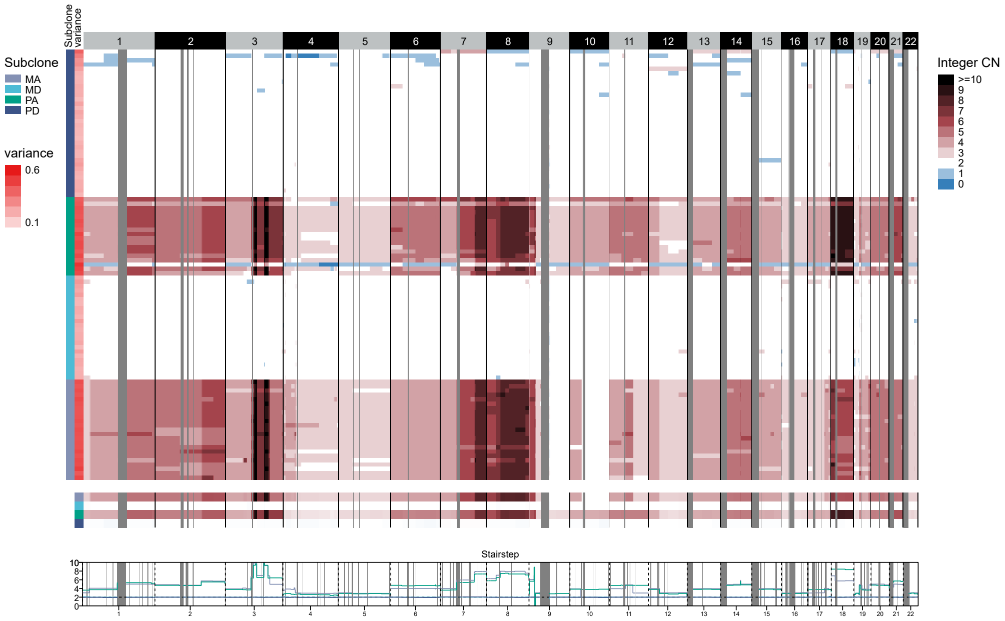


Figure S4: **The copy number profiles produced by SeCNV using hg38 reference.** For cells from T10 (A) and cells from T16 (B), SeCNV successfully reproduces the copy number profiles using hg38 reference.

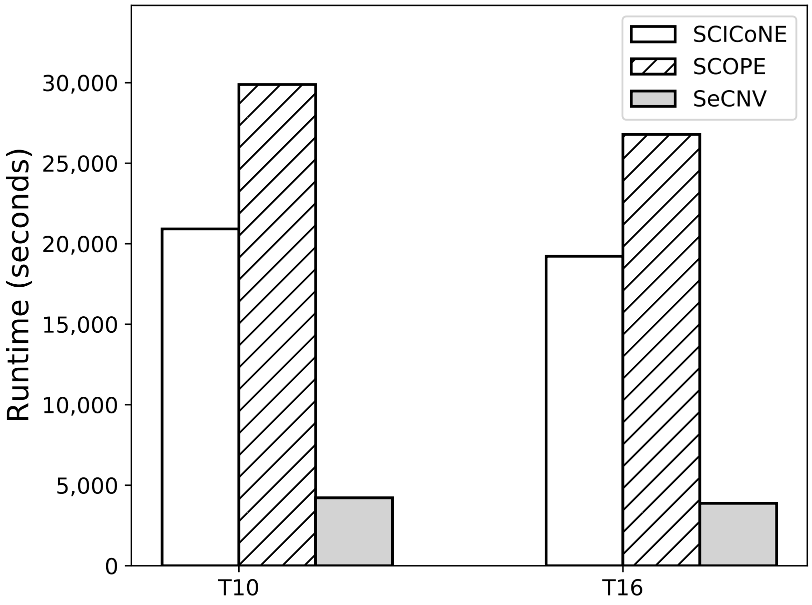


Figure S5: **Runtime of SCOPE and SeCNV on T10 and T16 datasets.** We compared the runtime of SCOPE and SeCNV for processing T10 and T16 datasets.

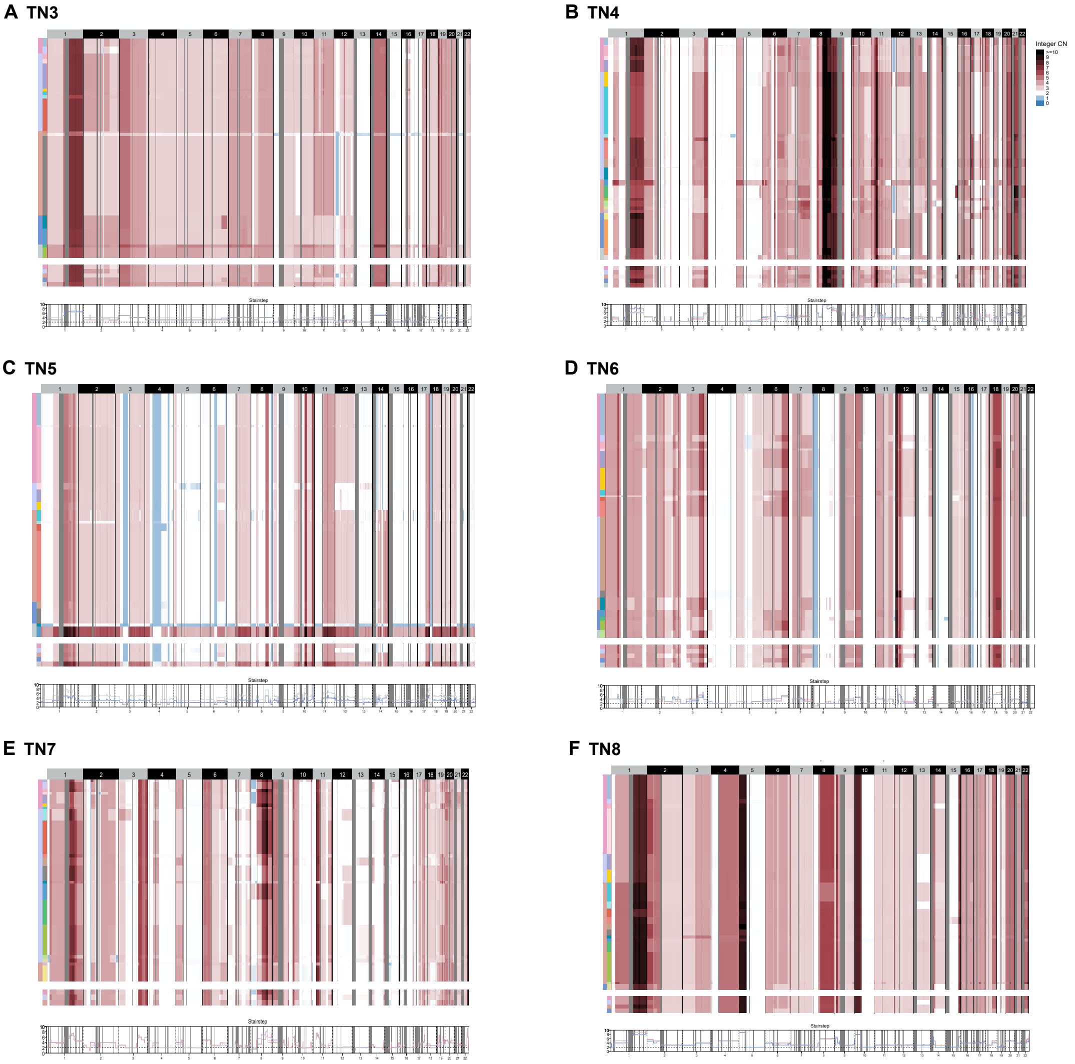


Figure S6: **The copy number profiles of TN3-TN8 estimated with SeCNV.** The two left columns show the superclones and subclones determined with copy number clustering.

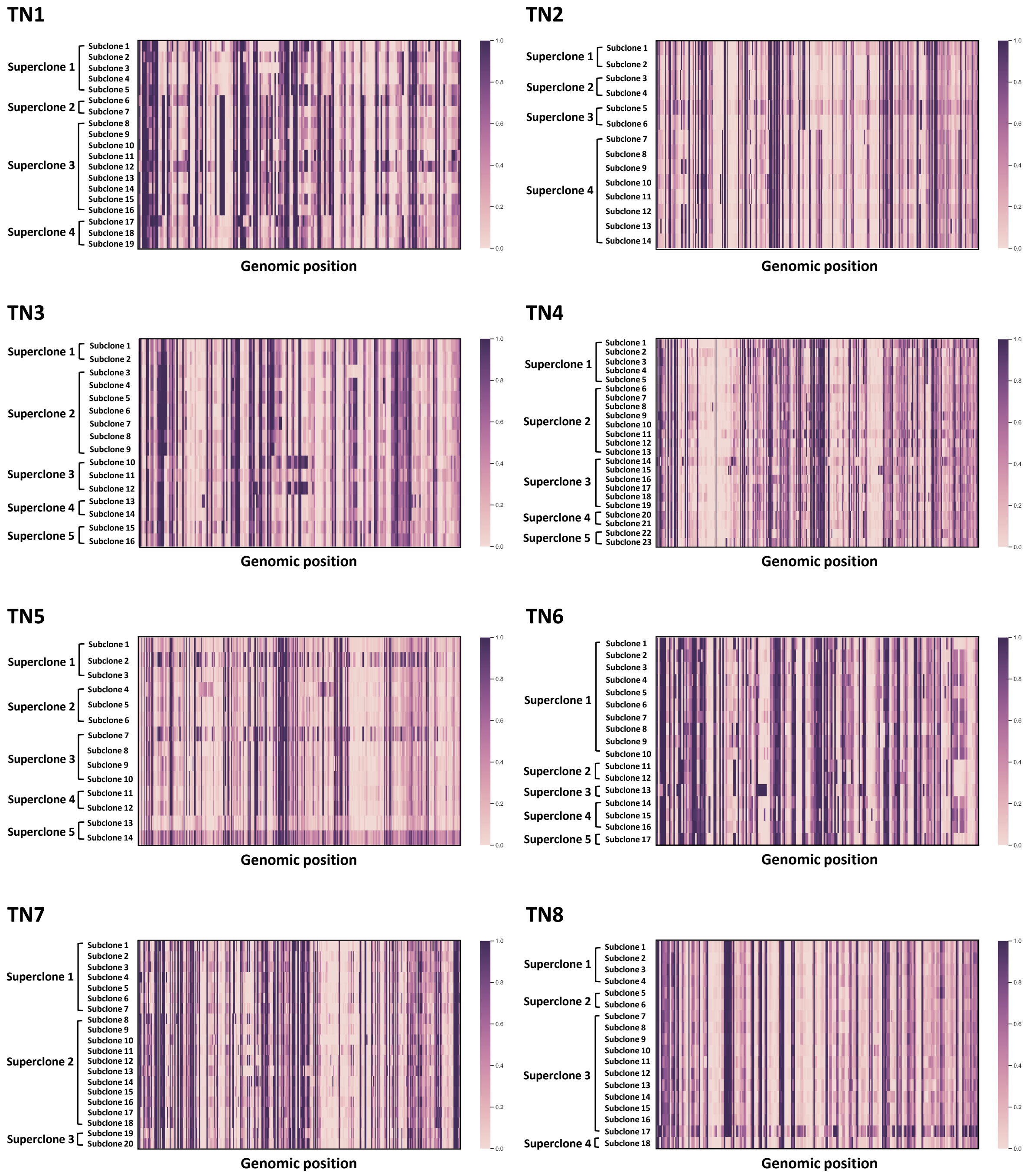


Figure S7: **The heatmaps of breakpoint occurrence probabilities for each subclone.** For subclones from TN1 to TN8, we calculated the breakpoint occurrence probabilities. We sorted the breakpoints according to their positions on the genome and the subclones according to the superclones they belong to.
